# Evolutionary dynamics of the vertebrate *Wnt* gene repertoire

**DOI:** 10.1101/2025.07.04.663182

**Authors:** Lily G. Fogg, Maxime Policarpo, Walter Salzburger

**Author notes:** Corresponding authors: Lily G. Fogg, Maxime Policarpo. These authors contributed equally (Lily G. Fogg, Maxime Policarpo).

## Abstract

The *Wnt* gene family plays a central role in vertebrate development, yet its evolutionary dynamics across lineages remain largely unexplored. Here, we present the most comprehensive analysis of *Wnt* gene evolution in vertebrates to date, leveraging a large-scale comparative genomics approach of 38,886 *Wnt* gene sequences mined from 1,961 species. We first investigate overall patterns of gene retention, duplication, and loss, and then focus specifically on the impact of whole-genome duplications (WGDs). We uncover striking variation in *Wnt* gene repertoire sizes, with ray-finned fishes (Actinopterygii) exhibiting the largest repertoires – even after excluding taxa with recent WGDs. Notably, we identify extreme expansions in polyploid cyprinids, including an octoploid hybrid harbouring 99 *Wnt* genes, the highest number observed. Unexpectedly elevated *Wnt* copy numbers in diploid species, such as the Antarctic lanternfish (*Electrona antarctica*) and the brook lamprey (*Lethenteron reissneri*), point to lineage-specific expansions with potential adaptive significance. Evolutionary rate analyses reveal that certain *Wnt* clades – especially *Wnt8* and *Wnt16* – exhibit elevated dN/dS ratios and high birth–death rates, indicative of repeated episodes of relaxed constraint or adaptive diversification. Contrary to our expectations, there was no relationship between developmental expression timing and evolutionary rates, suggesting pleiotropic regulation and functional redundancy of *Wnt* genes. Altogether, our findings reveal pervasive, lineage-specific remodelling of *Wnt* gene repertoires, shaped by both genome duplication history and divergent evolutionary trajectories. This work provides a high-resolution framework for understanding the molecular evolution of a key developmental toolkit and highlights candidate genes for future studies of vertebrate eco-morphological diversity.

## Introduction

The development and maintenance of animal form and function are regulated by signalling pathways that govern key processes, such as cell fate determination, tissue patterning, and organogenesis. Among these, the Wnt signalling pathway plays a pivotal role in development, influencing a wide array of biological processes including embryogenesis, stem cell maintenance, and the regulation of cell proliferation and differentiation (*1-3*). Given its central importance, dysregulation of the Wnt pathway is implicated in a variety of diseases, including cancer and developmental disorders (*4, 5*). The WNT ligands, key components of the Wnt signalling pathway, are secreted signalling proteins encoded by a family of genes that have been found in species across eukaryotes, from sponges to humans (*6, 7*).

In vertebrates, the *Wnt* gene family includes a large repertoire of genes that can be classified into at least 12 subfamilies based on their functional properties and sequence homology (*7*). Each gene clade is expressed at different points during development and plays distinct roles in developmental processes. Due to its critical role in development and cellular regulation, the Wnt signalling pathway is thought to be highly conserved across vertebrates (*8*). Furthermore, because morphological divergence increases over ontogeny, genes expressed at early and middle stages of development are thought to evolve under stronger purifying selective pressure than genes expressed late in development (*9*). However, these conclusions were drawn from studies focusing on a limited number of species or specific lineages. Despite the critical role of the *Wnt* genes in vertebrate biology, the evolutionary history and dynamics of the *Wnt* gene family and their link to expression patterns remain poorly understood beyond specific knowledge generated from a handful of model organisms.

Here, we conduct a comprehensive analysis of the *Wnt* gene family across a diversity of vertebrate species, utilizing a large-scale genomic approach to explore the evolutionary dynamics of these genes. By mining 1,961 vertebrate genomes, we characterized the diversification, gene gain and loss patterns, and lineage-specific expansions of the *Wnt* gene repertoire across all major vertebrate clades (ray-finned fishes, amphibians, reptiles, birds, and mammals). Furthermore, we investigated the potential associations between the *Wnt* gene repertoire and the developmental timing of gene expression and lineage-specific morphologies. Through this study, we provide a deeper understanding of the evolutionary history and functional diversification of the *Wnt* gene family across vertebrates and its implications for development and disease.

## Results and Discussion

### Vertebrate Wnt gene repertoire variation and expansion after whole-genome duplication

By extracting *Wnt* genes from 1,961 vertebrate species, we found that the number of *Wnt* genes per genome varied substantially across species and clades (Fig. 1; Fig. S1), reflecting both ancestral genome duplication events and lineage-specific evolutionary processes. Notably, while the loss of *Wnt* genes in a given species can arise from genome sequencing or assembly artifacts, our dense species sampling leads to an overall estimation of the number of *Wnt* copies across species reflecting true biological variations. This is supported by a strong phylogenetic signal (λ = 0.9, *p* < 2e-16) in copy number across the vertebrate phylogeny, *i.e,* closely related species tend to have similar numbers of *Wnt* genes, consistent with a pattern shaped by shared evolutionary history.

**Fig. 1.**
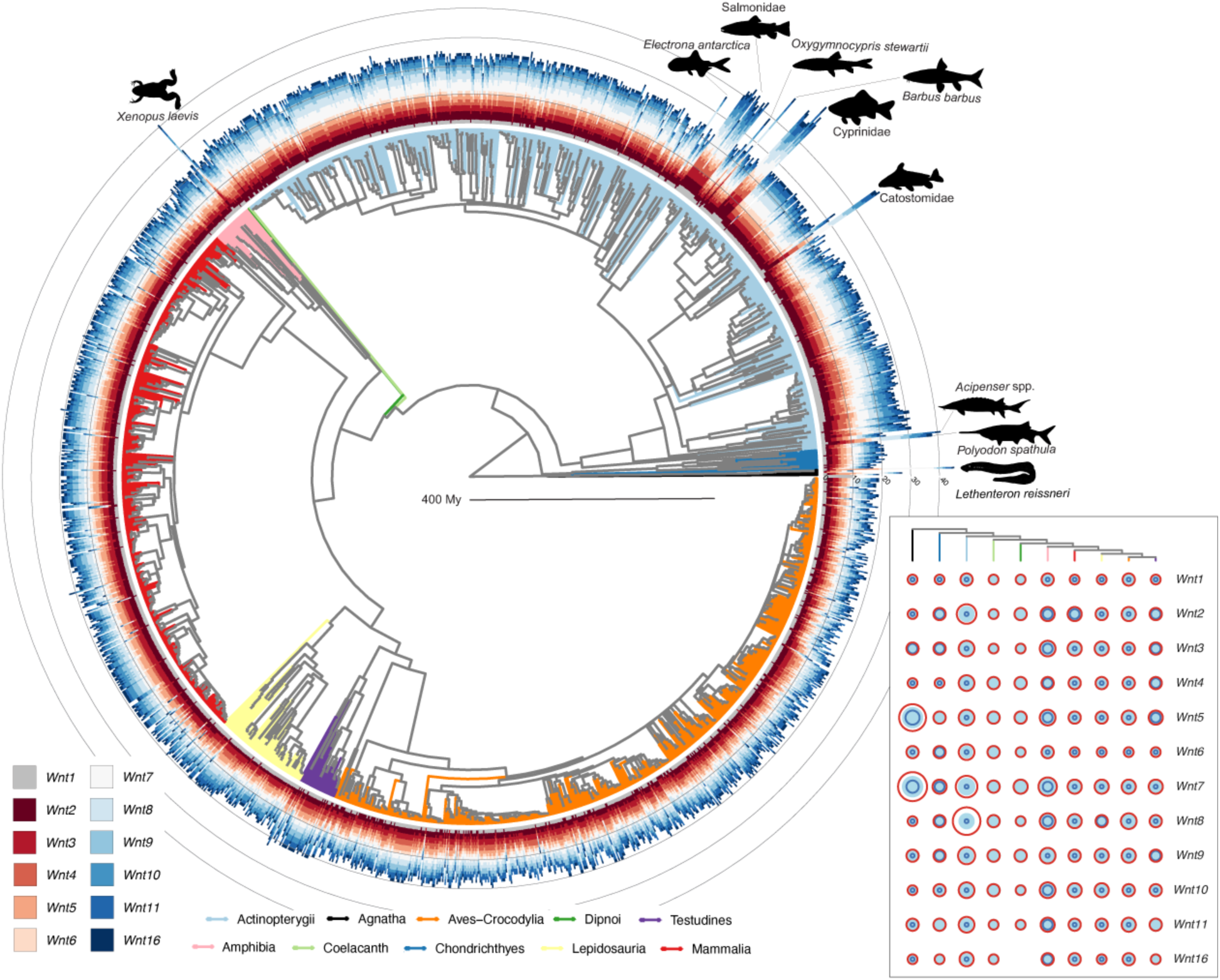
The *Wnt* gene repertoire in vertebrates. Phylogeny with 1898 vertebrate species, for which a genome assembly with more than 90% complete BUSCO genes was available. The branches are coloured by (sub)class. The number of *Wnt* genes per species are shown as bars, coloured as indicated in the lower left panel. Species with expanded *Wnt* gene repertoires are highlighted (indicated by black silhouettes). For the most part, these species have undergone additional whole-genome duplications, except for *Electrona antarctica* and *Lethenteron reissneri*. Inset plot is a visualisation of the size of the *Wnt* gene repertoire per gene clade per species class. Branches in phylogeny at the top are coloured by (sub)class. The minimum, mean and maximum copy numbers are represented by dark blue, light blue and red circles, respectively. Note that if there is no dark blue circle visible then the minimum is equal to the maximum. A phylogeny with full species names is available on the Figshare repository. Animal silhouettes were obtained from PhyloPic.org.

All *Wnt* subfamilies were retrieved in all vertebrate classes, with the exception of *Wnt16*, which could not be retrieved from the Dipnoi genome investigated here (*Neoceratodus forsteri*). Notably, Actinopterygii (ray-finned fishes) exhibited the largest *Wnt* gene repertoires, with a median of 24.6 *Wnt* genes per genome compared to 16.6 – 22 genes in the other vertebrate classes (Agnatha: 21.8; Chondrichthyes: 19.3; Coelacanth: 19; Dipnoi: 17; Amphibia: 20.9; Mammalia: 17.5; Lepidosauria: 16.9; Aves/Crocodilia: 16.7; Testudines: 19.6). The higher number of *Wnt* genes in ray-finned fish genomes is probably the result of a high retention of members of this gene family subsequent to the teleost whole genome duplication (WGD), and potentially linked to the extremely diverse morphologies in this class (*10*). Notably, this pattern persisted even after excluding non-diploid species, which have undergone more recent lineage-specific WGDs. The polyploid species, which had the highest *Wnt* copy numbers observed, included several linages in the Salmonidae, Cyprinidae, Catostomidae, and Acipenseridae families (Fig. 1). Particularly striking was the hybrid octoploid species *Carassius auratus × Cyprinus carpio*, which had 99 *Wnt* genes, followed by its tetraploid parental species, *C. auratus* and *C. carpio,* with 55 and 51 genes, respectively.

By leveraging these recent and independent WGD events in teleosts, we assessed whether certain *Wnt* subfamilies were preferentially retained. The average ratio of *Wnt* gene numbers in polyploid species compared to closely related diploid species ranged from 1.53 for *Wnt6* to 2.18 for *Wnt16* (Fig. S5). However, these differences were not statistically significant, suggesting either that there is no preferential retention of particular *Wnt* subfamilies following WGDs, or that these WGDs are too recent for sufficient post-duplication gene loss to reveal clear patterns of differential gene retention. Furthermore, while, as expected, we could observe significantly higher evolutionary rates of *Wnt* coding sequences in polyploid species (*p* = 4.699e-05, Fig. S5), we could not decipher if these accelerations were mainly driven by positive or relaxed selection, *i.e.,* driven by neo-functionalisation or non-functionalisation, also likely due to the relatively recent nature of these WGD events.

### Wnt gene repertoire expansion without additional whole-genome duplication

Among species without additional WGD events, elevated *Wnt* gene numbers were found in the Antarctic lanternfish *Electrona antarctica* (36 genes), the Asiatic brook lamprey *Lethenteron reissneri* (45 genes) and most species in the Eloposteoglossocephala (*11*) (range: 24 – 31 genes; mean: 28 genes) (Fig. 1). This finding suggests either localized gene family expansion or possible assembly artifacts (*12*).

To investigate these unusual cases further, we examined the genomic characteristics of *Wnt* genes in *E. antarctica* and *L. reissneri*. In *E. antarctica*, the expansion was driven predominantly by remarkable duplications of *Wnt8a.1*, with 14 copies – more than in any other species examined (Fig. S2). Genomic coverage levels for these paralogs were similar to other *Wnt* genes and to BUSCO reference genes, indicating that these duplicates are unlikely to be assembly artifacts (Fig. S2). However, experimental validation (*e.g.,* via Sanger sequencing) would be required to confirm their authenticity. Evolutionarily, the *Wnt8a.1* genes in *E. antarctica* exhibited elevated dN/dS ratios relative to both *Wnt8a.1* orthologs in other species and to other *Wnt* genes, suggesting a history of relaxed purifying selection or possible positive selection (Fig. S2). The relatively compact size of *Wnt8a* in fish (in *E. antarctica*, *Wnt8a* is approximately 1.5 kb compared to, for example, *Wnt4a* at >20 kb), together with its genomic architecture and known bicistronic structure (*13*), may facilitate tandem duplication, potentially contributing to its lineage-specific expansion.

In *L. reissneri*, the elevated *Wnt* copy number was due to expansions in *Wnt5a* and *Wnt7bb*, with 9 and 13 copies respectively – again the highest copy numbers in these gene clades among all species examined (Fig. S3). Genomic coverage for these paralogs was lower than BUSCO genes but consistent with other *Wnt* genes in this genome (Fig. S3), leaving their authenticity inconclusive. These patterns may reflect true expansions or potential assembly or annotation issues, which are not uncommon in lamprey genomes due to their high repeat content and fragmented assemblies (*14*). Nonetheless, if genuine, these expansions point to ongoing *Wnt* diversification even in early-diverging vertebrate lineages such as Agnatha.

In Eloposteoglossocephala, which includes Osteoglossomorpha (*e.g.,* Arapaima and elephantfishes) and Elopomorpha (eels and tarpons), the increased size of the *Wnt* gene repertoire was due to elevated copy number across several *Wnt* clades, specifically *Wnt1, Wnt2, Wnt4, Wnt6, Wnt10, Wnt11,* and *Wnt16*. These slightly more elevated copy numbers were consistently observed across species in the clade (Fig. 1). The reasons for this pattern may arise from higher retention of duplicates from the teleost WGD compared to Clupeocephalans. Additionally, ecological or morphological factors unique to Eloposteoglossocephala may have favoured the maintenance of extra *Wnt* copies. Indeed, species in this clade exhibit a wide range of life history strategies, such as long-distance catadromous migrations, air-breathing adaptations, and reproductive strategies involving complex parental care (*15-17*). Thus, the elevated *Wnt* copy numbers in this clade may reflect adaptive pressures related to the group’s distinctive evolutionary and ecological contexts, independent of whole-genome duplication.

### Selective pressures and birth-death dynamics of Wnt genes

To assess the evolutionary dynamics of *Wnt* genes, we first analysed codon-level substitution rates (dN/dS) across gene clades and across vertebrate classes and then used gene tree – species tree reconciliation methods to compute duplication and loss rates (per gene per million years) for each subfamily and each vertebrate class. While the overall distributions of *Wnt* dN/dS values were similar between vertebrate classes (Fig. 2; Fig. S4), there were striking differences in terms of duplication rates. The most dynamic *Wnt* gene repertoires were found in Actinopterygii and Mammalia, with significantly higher birth rates than in the other classes (Fig. 3a), as well as higher death rates than in most classes (Fig. 3b). In fishes, the expanded and highly dynamic *Wnt* repertoires might have contributed to the exceptionally diverse morphological features of this group, along with their diversification and adaptation to diverse aquatic niches. In both ray-finned fishes and mammals, gene clades with the highest birth rates included *Wnt8*, *Wnt10*, and *Wnt16* (Fig. 3c and 3d). Notably, this pattern persisted even when polyploid fish clades (which inflate birth rate estimates) were excluded. Furthermore, in all vertebrate classes except in Agnatha, we observed that, with a mean dN/dS of 0.237, *Wnt16* genes were evolving faster than the other subfamilies, which had mean dN/dS ranging from 0.03 for *Wnt1* to 0.16 for *Wnt10* (Fig. 3e).

**Fig. 2.**
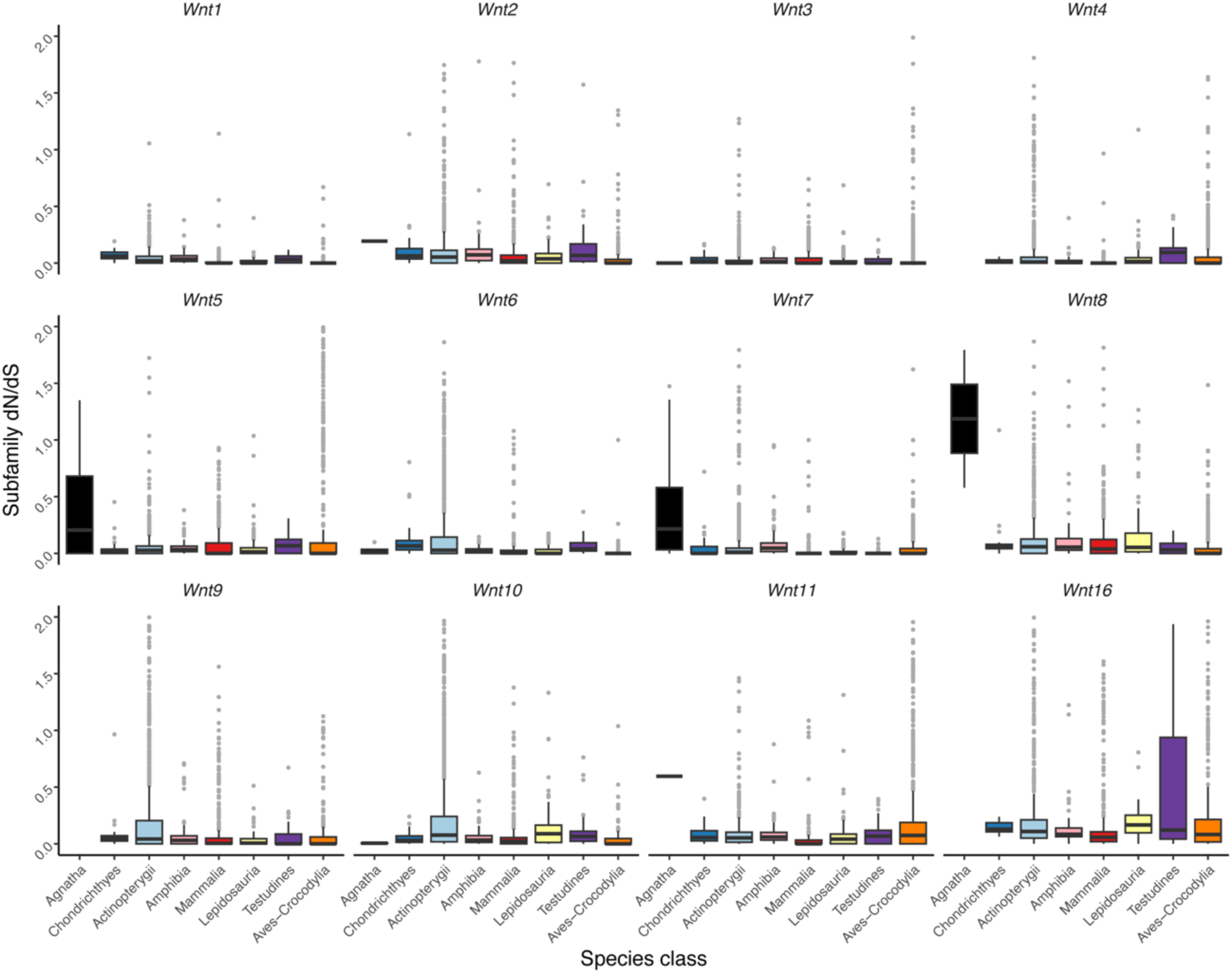
dN/dS of *Wnt* genes across vertebrate classes. For each vertebrate (sub)class (coloured as in Fig. 1), the distributions of the dN/dS values of the different *Wnt* genes are shown as boxplots (first quartile −1.5 interquartile range; first quartile; mean; third quartile; third quartile +1.5 interquartile range; dots represent outliers) for all vertebrate species for which a genome assembly with more than 90% complete BUSCO genes was available. Samples sizes for each vertebrate (sub)class can be retrieved from Supplementary Table 1.

**Fig. 3.**
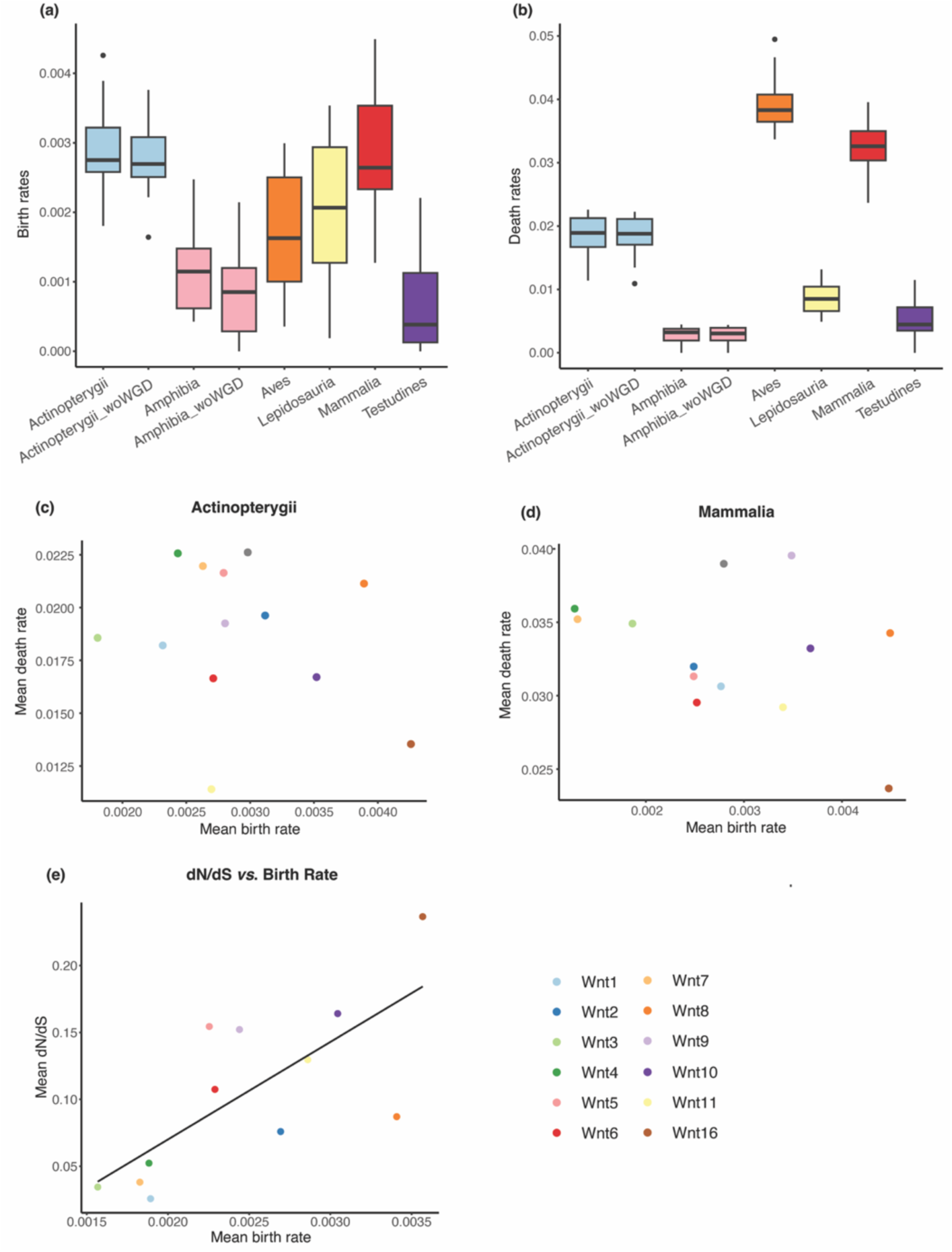
Birth and death rates of *Wnt* genes across vertebrate classes. Birth **(a)** and death **(b)** rates of all *Wnt* genes per vertebrate class (coloured as in Fig. 1) shown as boxplots (first quartile −1.5 interquartile range; first quartile; mean; third quartile; third quartile +1.5 interquartile range; dots represent outliers). Note that the classes containing representatives with additional whole-genome duplications (WGDs) have been split into species with (Actinopterygii and Amphibia) and without (Actinopterygii_woWGD and Amphibia_woWGD) these additional WGDs. **(c-d)** Mean birth *vs.* death rates per *Wnt* clade for the classes with the highest birth rates: Actinopterygii (c) and Mammalia (d). **(e)** Relationship between mean dN/dS and mean birth rate per *Wnt* clade. Black line shows a linear regression. Note that the colour legend for the different *Wnt* clades represented in panels c-e is located in the bottom right of the figure.

The elevated duplication rates in these *Wnt* gene clades suggest that the biological functions associated with *Wnt8*, *Wnt10*, and *Wnt16* may be subject to increased lineage-specific diversification. For example, the dynamics observed in *Wnt8* could reflect its central role in embryonic development – particularly axis formation, neural patterning, and somitogenesis (*13, 18*) – where shifts in expression timing or regulatory architecture may promote developmental innovation. Similarly, the high turnover of *Wnt10* may relate to its roles in epithelial-mesenchymal interactions, skin appendage and hair follicle development (*19, 20*), and bone morphogenesis (*21*), processes that are often tightly linked to lineage-specific morphological traits. Finally, *Wnt16* is implicated in diverse biological processes, including bone homeostasis in mammals (*22*), as well as hematopoietic stem cell development in zebrafish (*23*), indicating its functional versatility across vertebrates.

In contrast, certain *Wnt* subfamilies exhibited remarkably low duplication rates across vertebrate classes, suggesting strong evolutionary constraint. *Wnt1, Wnt3, Wnt4,* and *Wnt7* were the least dynamic gene clades, with low mean dN/dS values and birth rates (Fig. 2; Fig. 3e). This evolutionary conservation suggests that the functions of these genes are highly constrained and likely important across diverse lineages. Indeed, these gene subfamilies are involved in critical and conserved developmental processes. For instance, *Wnt1*, *Wnt3* and *Wnt7* all play central roles in the development of the nervous system (*24-31*), while *Wnt4* is crucial for gonad development and sexual maturation (*32, 33*). The evolutionary stability of these genes implies that functional indispensability may limit the tolerance for duplication and mutation, leading to strong purifying selection that maintains their structure and function across vertebrates.

Finally, we observed a positive correlation between the mean birth rate of *Wnt* subfamilies and their mean evolutionary rate (linear model: R^2^ = 0.54; *p* = 0.007; Fig. 3e), showing that when genes are being duplicated, the evolutionary rates of their coding sequences increase, consistent with their neo-, sub- or non-functionalisation. Notably, the mean birth rate of *Wnt16* was also the highest (0.004), followed by *Wnt8* (0.0034) and *Wnt10* (0.003).

### Pleiotropy, developmental constraint, and lineage-specific selection in Wnt gene evolution

To investigate whether developmental expression patterns influence the evolutionary dynamics of *Wnt* genes, we analyzed three well-characterized model organisms representing major vertebrate clades: *Danio rerio* (Actinopterygii), *Xenopus laevis* (Amphibia), and *Mus musculus* (Mammalia). Specifically, we tested the hypothesis that *Wnt* genes expressed earlier in development would exhibit lower evolutionary rates, as early-expressed genes are often subject to stronger purifying selection due to their involvement in foundational developmental processes (*34-36*).

Contrary to expectations, no significant correlation was detected between the timing of developmental expression and dN/dS values across species (Fig. S7-9). This lack of association may reflect the complex and often stage-specific roles of *Wnt* genes during development. Wnt signalling is known to act repeatedly across embryogenesis and later life stages, with many genes playing roles in multiple tissues and developmental contexts. As a result, even genes with early embryonic expression may be subject to divergent evolutionary pressures across other stages or tissues. Additionally, the high degree of pleiotropy observed in *Wnt* genes may buffer their evolutionary rates against simple temporal constraints. In such cases, compensatory mechanisms, redundancy within the *Wnt* gene family, or dosage sensitivity may dilute the relationship between expression timing and sequence constraint. Moreover, available developmental expression datasets may not fully capture the spatial-temporal complexity of *Wnt* gene regulation, especially for genes with dynamic or tissue-specific expression.

While global patterns of evolutionary constraint in the *Wnt* gene family did not correlate with developmental timing, we tested if we could detect signatures of shifts in selective pressure during specific evolutionary transitions. Given the central role of Wnt signalling in appendage development, we explored the relationship between *Wnt* gene evolution and the loss of limbs, which happened several times independently in amphibian and lepidosaurians (six independent times when considering species included in this study; Fig. 4a).

**Fig. 4.**
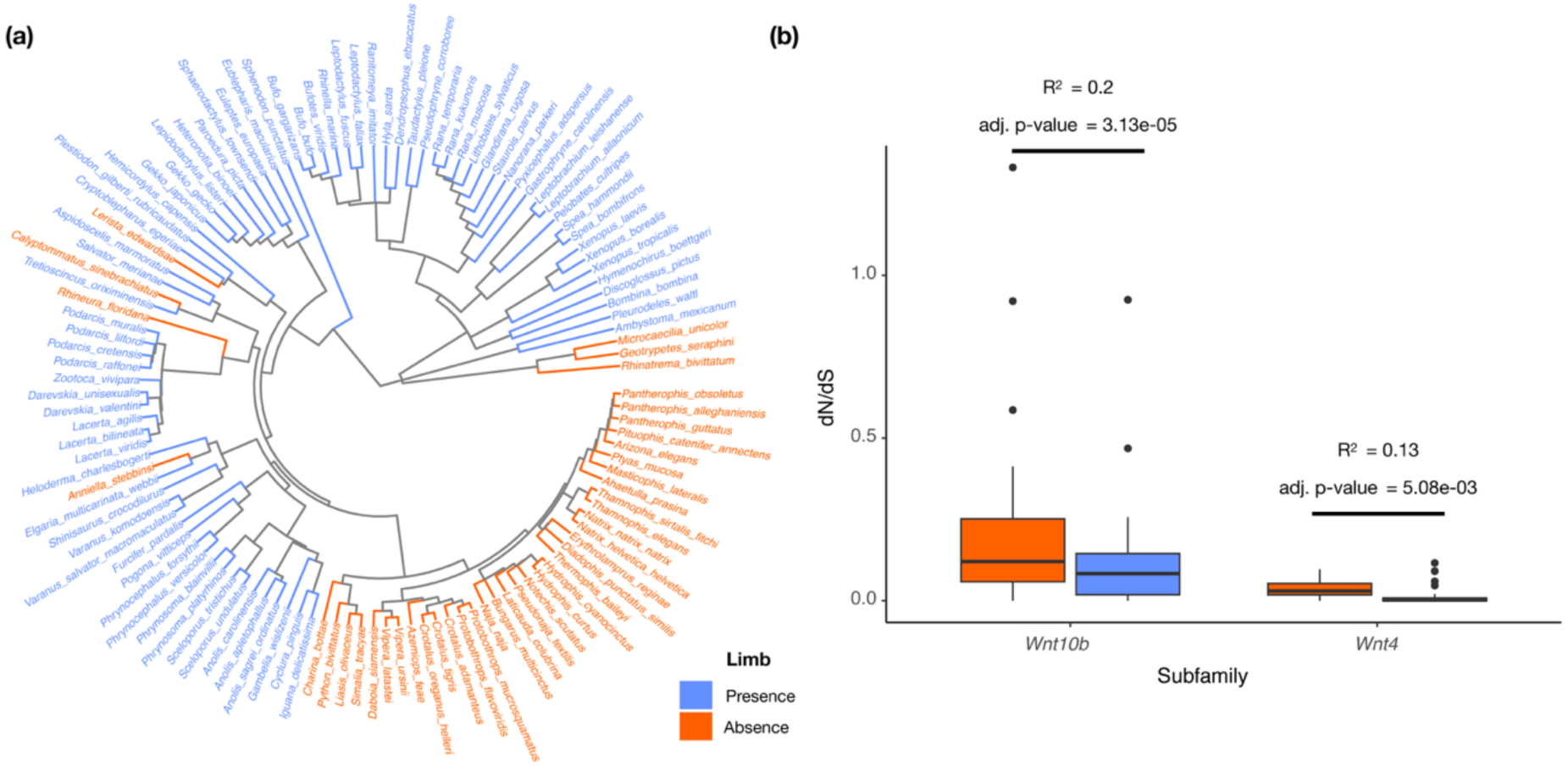
The dynamics of *Wnt* genes in lineages with limb loss. **(a)** Phylogeny of diploid amphibian and lepidosaurian species. Species with limbs are displayed in blue while species which have undergone limb loss are displayed orange. **(b)** Relationship between the dN/dS values of *Wnt10b* and *Wnt4* and the presence or absence of limbs. pGLS R^2^ and adjusted *p* values (Bonferroni correction) are indicated.

Strikingly, we found that *Wnt10b* and *Wnt4* exhibited significantly elevated dN/dS ratios in lineages that have undergone limb loss (pGLS R² values of 0.20 and 0.13, with corresponding adjusted *p* values of 3.13e-05 and 0.13, respectively; Fig. 4b; Supplementary Table 1), suggesting a relaxation of purifying selection or the onset of positive selection following the loss of limbs. These findings are consistent with the known roles of both genes in appendage development. *Wnt4* has been implicated in early limb bud formation and mesenchymal cell proliferation (*37, 38*), while *Wnt10b* is involved in bone development and outgrowth and patterning of the limb (*21, 39, 40*). The elevated dN/dS values in limbless taxa indicate that these genes may have experienced reduced functional constraints or adaptive shifts in regulatory or developmental roles following the loss of limbs.

### Conclusion

In this study, we have assembled a comprehensive *Wnt* gene atlas from nearly 2000 vertebrate genomes, providing a valuable resource for future comparative and functional studies in non-model organisms. This study reveals widespread variation in vertebrate *Wnt* genes, shaped by both whole-genome duplications and lineage-specific expansions. Teleosts, particularly polyploid species, show the most extensive *Wnt* gene repertoires, but we also identify unexpected gene expansions in species without recent WGDs, indicating localized duplication events.

Gene duplication and evolutionary rate analyses highlight *Wnt8, Wnt10,* and *Wnt16* as particularly dynamic subfamilies, potentially contributing to lineage-specific morphological innovation. In contrast, the strong conservation of *Wnt1, Wnt3, Wnt4,* and *Wnt7* underscores their essential developmental roles. The positive link between duplication rate and sequence evolution supports models of post-duplication divergence.

Although developmental expression timing did not predict sequence constraint, shifts in selective pressure following limb loss in amphibians and lepidosaurs, especially in *Wnt4* and *Wnt10b*, indicate that changes in the evolutionary rates of genes can accompany major morphological transitions. Together, our findings highlight how genome architecture, developmental function, and ecological context interact to shape the evolution of a key signalling pathway in vertebrates.

## Materials and Methods

### Genome Data

All genome assemblies available on the National Center for Biotechnology Information (NCBI) database (*41*) on 6^th^ November 2023 were downloaded using genome_updater and the following options: -T “7742” (restricts the genome search to vertebrate species), -d “refseq,genbank” (searches both the RefSeq and Genbank databases), and -A 1 (retains only one genome assembly per species). A total of 3533 vertebrate genomes were initially downloaded, of which 169 were excluded as the assemblies were described as “partial”, leaving a total of 3360 vertebrate genomes (Agnatha: 8; Chondrichthyes: 26; Actinopterygii: 1325; Dipnoi: 2; Coelacanth: 1; Amphibia: 85; Mammalia: 776; Lepidosauria: 122; Testudines: 36; Crocodilia: 5; Aves: 974). Alternative scaffolds were removed from the *Homo sapiens* (<u>GCA_000001405.40</u>) and *Danio rerio* (<u>GCA_000002035.6</u>) genomes.

### Genome completeness assessment

The completeness of the vertebrate genomes used for this study was assessed using a reimplementation of BUSCO, known as compleasm v0.2.4 (*12, 42*), using the vertebrata odb10 database, except for three extremely large genomes (Dipnoi: *Protopterus annectens* and *Neoceratodus forsteri*; Amphibia: *Ambystoma mexicanum)*, for which BUSCO results were retrieved from previous studies (*43-45*). Since it is expected that genomes with a large proportion of missing BUSCO genes will produce biased estimates for the number of *Wnt* genes, we only selected high-quality genome assemblies on the basis of a high BUSCO score threshold: 90% complete BUSCO genes. In jawed vertebrates, 1955 genome assemblies contained at least 90% complete BUSCO genes. Consistent with previous studies, and likely due to the lack of a suitable BUSCO gene dataset for agnathan species, the eight agnathan genome assemblies had low BUSCO scores (between 17.1 and 76.4%) (*46*). Thus, the five highest agnathan genomes (with BUSCO scores above 60%) were retained in our analyses. Similarly, there were only two genome assemblies available for Dipnoi (*Neoceratodus forsteri* and *Protopterus annectens*), and the best of these two was retained (*N. forsteri*, 83.1% complete) to represent this class. Finally, one lepidosaur species (*Sceloporus occidentalis*) was removed from the analyses, as this genome is contaminated with amphibian DNA (*46*).

### Vertebrate species tree

The phylogenetic context of the species retained after BUSCO filtering was estimated by reconstructing a vertebrate species tree as described in (*46*). All subspecies were discarded to avoid redundancy in the species representation, leaving 1898 species for the phylogeny. Firstly, single-copy orthologs were identified by running BUSCO v5.7.1 (*12*) on each genome assembly using the closest available database (*Actinopterygii odb10 database*: Actinopterygii; *Aves odb10*: Aves; *Mammalia odb10*: Mammalia; *Tetrapoda odb10*: Amphibia, Crocodylia, Lepidosauria, and Testudines; *Vertebrata odb10*: Agnatha, Chondrichthyes, Coelacanth, and Dipnoi). The genome assemblies of two species had to be split prior to running BUSCO due to their large size (*Rana muscosa* and *Ambystoma mexicanum*). Single-copy orthologs present in at least half of the species were aligned using MUSCLE v5.1 (*47*) and alignments trimmed using trimAl v1.4.1 (*48*). Maximum likelihood gene trees were constructed for each ortholog using IQ-TREE v2.0 using automated model selection by ModelFinder (*49*). We then inferred an unrooted species tree using ASTRAL v5.7.8 (*50*) and the resulting tree was rooted and dated using the least squared dating method (*51*) implemented in IQ-TREE v2.0 (using calibration dates, extracted from TimeTree.org (*52*), available in Supplementary Table 1). A vertebrate species tree was generated by merging these species tree, using the R package “ape” (*53*) and the function “bind.tree”.

### Wnt gene mining

First, we built a *Wnt* database by extracting annotated *Wnt* genes (GenBank and RefSeq databases) from 28 vertebrate species. Genes annotated as partial or as pseudogenes were discarded, and we verified that retained genes were best matching to *Wnt* sequences by performing a BLASTP against the UniProt database. For each genome included in this study, we first performed a TBLASTN using this *Wnt* database as query, the genome as target and an e-value of 1e-3. We then extracted nonoverlapping tblastn hit regions using SAMtools, and these regions were extended by 100,000 bp upstream and downstream. Again, after this extension, overlapping regions were merged. Potential *Wnt* genes were predicted on these regions using miniprot, using the *Wnt* database as query. We discarded miniprot predictions for which the number of exons was lower than the number of exons inferred from tblastn results (*i.e.,* number of non-overlapping tblastn hit in this gene region), as this was indicative that miniprot likely merged two nearby *Wnt* genes. Miniprot results were further filtered to keep only predictions which lengths were higher than 95% of the query (*i.e.,* higher than 95% of a *Wnt* genes from our *Wnt* database). If two or more predictions on the same genomic region met these criteria, we kept the one that spanned the shortest genomic region. We repeated this filtering operation by decreasing the threshold length, until no miniprot prediction was found any more. Each miniprot predictions were translated to a protein sequence using EMBOSS transeq and this protein was used as query in a blastp against the UniProt database. Only predictions best-matching to a *Wnt* gene were retained. If a prediction contained one or more frameshifts, these were discarded before the translation. Retained miniprot predicted sequences were then classified as complete genes if they had a length higher or equal to 75% of their best blastp mach against the *Wnt* database, as pseudogenes if they contained one or more loss-of-function mutations (premature stop codon or frameshift), or as incompletes if no loss-of-function mutation was retrieved, and if their length was shorter than 75% of their best blastp mach against the *Wnt* database. We also classified predictions containing stretches of ambiguous (“N”) nucleotides as incomplete genes.

### Assessment of gene classification bias and species filtering criteria

To ensure that genes were not disproportionately assigned to the incomplete or pseudogene categories, correlations were assessed between the number of complete genes and the number of incomplete genes or pseudogenes. These analyses were conducted both across all species and within a subset of species which had a lower number of complete *Wnt* genes, *i.e.,* fewer than 12 (Fig. S6). A very weak negative correlation was identified between the number of complete genes and pseudogenes, but not between complete and incomplete genes, when all species were considered. However, this correlation was not observed when analyses were restricted to species with low numbers of complete *Wnt* genes. These findings suggest that contracted *Wnt* repertoires in these species are unlikely to result solely from pseudogenisation or misclassification by the gene mining pipeline. Nevertheless, to minimize potential bias arising from genome assembly or annotation quality, species with exceptionally low numbers of complete *Wnt* genes (fewer than six) were excluded from downstream analyses.

### Wnt gene trees and gene verification

Gene trees were reconstructed using similar methods for each of the following groups: 1) all *Wnt* sequences mined, 2) per *Wnt* gene subfamily, and 3) per *Wnt* gene subfamily and species class. Firstly, given the large number of gene sequences, a template alignment was first generated by aligning 30 random sequences from the dataset using MUSCLE v5.1 (*47*) and the final alignment was generated by adding all other sequences to the template using the option “-add” in MAFFT v7.467 (*54*). For the per-gene clade trees, 10 randomly selected sequences from another gene clade were also added as outgroups. The final alignment was trimmed using trimAl v1.4.1 (*48*) and maximum likelihood gene trees were generated using IQ-TREE v2.0 (*49*) with the optimal model determined by ModelFinder (*55*). Trees were rooted in iTOL (*56*) and visualised using Taxonium (*57*) and the R package ggtree (*58*).

Following initial gene tree construction, the *Wnt* gene sequences were manually verified. Firstly, visual inspection was used to identify sequences : 1) causing disparity between the topology of the gene tree and the vertebrate species tree, and 2) sequences with long branches which could represent erroneous predictions and/or distort the tree topology (*59*). All questionable sequences were manually re-predicted using a slower but more accurate exon prediction program, known as EXONERATE v2.2.0 (*60*). If the EXONERATE prediction disagreed with the prediction originally generated by miniprot, the former was used and the gene was accordingly (re-)classified as “complete”, “pseudogene”, or “incomplete”. This process was repeated iteratively, resulting in the manual verification of 995 genes. The final dataset comprised 38,886 complete *Wnt* gene sequences. We computed the phylogenetic signal of the total number of *Wnt* genes using the phylosig function implemented in phytools (*61*), with the lambda method.

### Sequence evolution analyses

Branch rates of sequence evolution (*i.e.*, ω or dN/dS ratio) were estimated from per-gene clade, per-species class alignments and phylogenies generated as described in the “*Wnt gene trees and gene verification”* section. To obtain maximum likelihood ω ratios per branch, we fitted the Muse Gaut (+ GTR) model of codon substitution (*62*), implemented in HyPhy (*63*). Branches with a lack (dS < 0.01) or a saturation of substitutions (dS or dN > 1) were discarded.

### Birth-death rate calculations

The per-gene clade, per-species class phylogenies were used to reconstruct the evolutionary dynamics of the *Wnt* gene family. Firstly, nodes with low bootstrap values (<95%) were collapsed into polytomies using the R package ape (*53*). TreeRecs (*64*) was used to find the best root per gene tree, and NOTUNG v2.9.1.5 (*65*) was used with the phylogenomics option to reconcile the rooted gene tree with the species tree for each gene clade and species class. From the reconciliation results, we extracted the number of duplications, losses, and gene copy number for each internal node. We then computed birth (λ) and death rates (μ) per gene per million years, from duplication and losses numbers, following the approach of Niimura *et al. (66)*. Branches smaller than two million years were excluded (*67*). Rates were calculated for each gene clade within each species class, and mean birth and death rates were computed by averaging birth and death rates across all branches.

### Developmental timing of Wnt gene expression and correlation with evolutionary rates

To investigate the relationship between developmental timing of *Wnt* gene expression and rates of molecular evolution, transcriptomic and sequence data were analyzed for three model species: *Danio rerio, Xenopus laevis,* and *Mus musculus*. For each species, a developmental time-course of gene expression was used to determine the timing of *Wnt* gene activity across embryogenesis. Expression data for *D. rerio* were obtained from the EMBL-EBI Expression Atlas (https://www.ebi.ac.uk/gxa/experiments/E-ERAD-475) and were based on an RNA-seq dataset published by White *et al.* (*68*). The *X. laevis* data were downloaded from Xenbase (https://www.xenbase.org) and were based on an RNA-seq dataset published by Session *et al. (69)*. The data for *M. musculus* were obtained from the MGI-Mouse Gene Expression Database (GXD; https://www.informatics.jax.org/expression.shtml).

For each gene, TPM values were used to define two key developmental time points: (i) peak expression, the stage with the highest TPM value; and (ii) earliest expression, the first stage with TPM > 0. Developmental stages were filtered to retain similar stages across species using on morphological criteria broadly conserved across vertebrates (*e.g.,* zygote, gastrulation, somitogenesis, and pharyngula stages). Each stage was assigned a numerical rank to permit regression-based analysis of expression timing as a continuous variable.

For each species, expression data were integrated with a curated set of *Wnt* gene sequences mined in this study. Publicly annotated transcripts from Ensembl (*70*) were matched to mined transcripts using phylogenetic inference, ensuring orthology and accurate mapping between expression profiles and evolutionary rate data. Only *Wnt* genes with both reliable expression classification and dN/dS estimates were included (see section on *Sequence evolution analyses* for details of dN/dS filtering).

To assess whether the timing of gene expression is predictive of evolutionary constraint, dN/dS values were regressed against the numerical ranks of peak and earliest expression stages using generalized least squares (GLS) models in R. This allowed for species-specific tests of the hypothesis that genes expressed earlier in development are subject to stronger purifying selection.

### Retention of Wnt genes following whole-genome duplications in teleosts

Across teleost species included in our study, there were six independent recent whole genome duplications: one in Salmoniformes, four in Cyprinidae, and one in Catostomidae. For each of these polyploid lineages, we computed the mean number of genes per *across polyploid* species, for each *Wnt* subfamily. In the same way, we computed the mean number of gene per species and per *Wnt* subfamily for close diploid species (species included available in Supplementary Table 1). Then, for each WGD lineage, we could compute the ratio between the with these mean number of *Wnt* genes and the mean number of *Wnt* genes in close diploid species. We tested if the computed ratios were significantly different between *Wnt* subfamilies using pairwise Wilcoxon tests (Supplementary Table 1). The *p* values were adjusted with the Bonferonni correction. For the same set of species, we then used RELAX, implemented in HyPhy, to test if branches corresponding to *Wnt* genes of polyploid species were evolving under intensified or relaxed selection. We also tested, as control, branches corresponding to *Wnt* genes of close diploid species. A branch was considered to evolve under relaxed selection if the RELAX *p* value was < 0.05 and the K parameter <= 1 or under intensified selection if the *p* value was < 0.05 and the K parameter > 1.

### Correlation between Wnt selective pressure and loss of limbs

For each *Wnt* gene retrieved in lepidosaurian and amphibian, we performed a pGLS using the R package “caper” (*71*) with, as response, the dN/dS and, as predictor, the presence or absence of limbs. Only *Wnt* genes retrieved in at-least three independent clades with limb losses were retained. Resulting *p* values were corrected using the Bonferroni method.

## Acknowledgements

We would like to thank the members of the Salzburger lab for valuable suggestions and comments on this study. All calculations were performed at sciCORE (http://scicore.unibas.ch/), the center of scientific computing at University of Basel [with support by the SIB (Swiss Institute of Bioinformatics)].

## Data Availability

Custom analysis scripts are provided on GitHub (https://github.com/MaximePolicarpo/WNT_vertebrates). All other data are available via Figshare or are provided in the main manuscript or Supplemental Information.

