## Supplementary Information for "Evolutionary dynamics of the vertebrate *Wnt* gene repertoire"

[#]These authors contributed equally

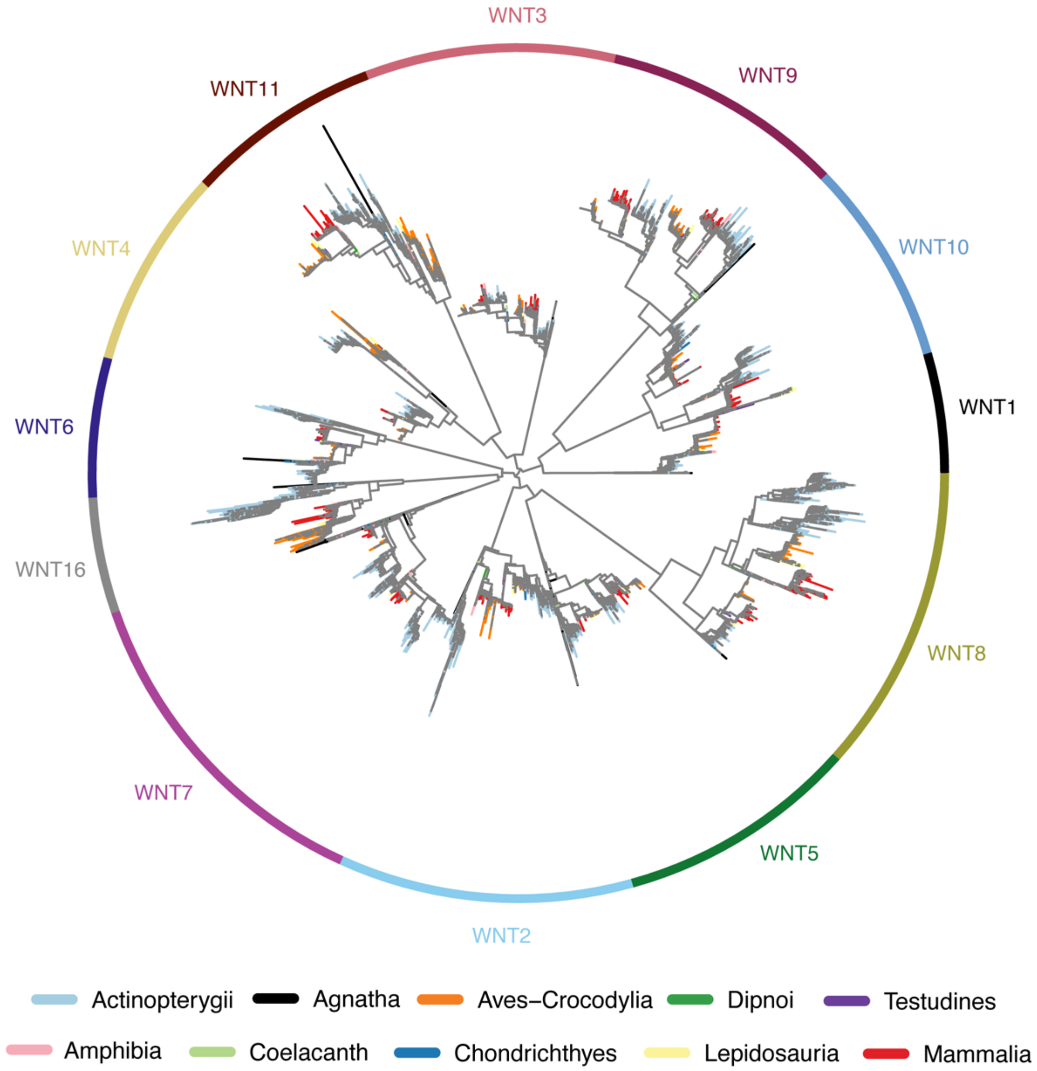


**Fig. S1. Vertebrate *Wnt* gene tree.** Gene tree containing all *Wnt* sequences mined in this study. Branches are coloured by species class and enclosing circle is coloured by *Wnt* gene subfamily.

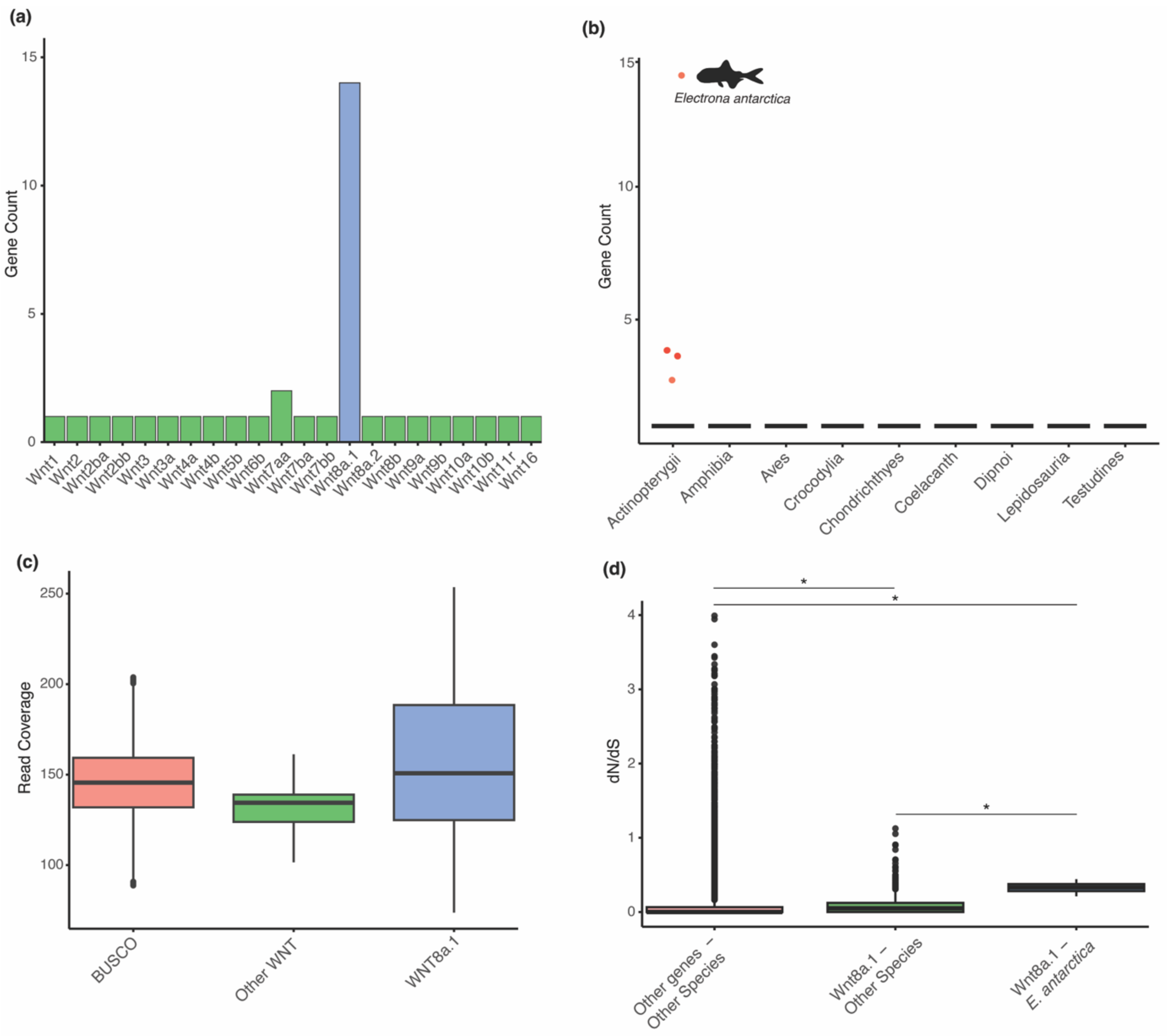


**Fig. S2. *Electrona antarctica Wnt8a.1* expansion. (a)** Gene copy number per *Wnt* subfamily in *E. antarctica* showing that the expansion of *Wnt* genes in this species is mostly due to *Wnt8a.1*. **(b)** The distribution of *Wnt8a.1* genes across all vertebrate species analysed, split by class. The species with the highest copy number of this gene (*E. antarctica*) is labelled. **(c)** A comparison of raw genomic read coverage between BUSCO genes, other *Wnt* genes (excluding *Wnt8a.1* copies), and only the *Wnt8a.1* genes in *E. antarctica*. **(d)** A comparison of omega ratios (dN/dS) across all species and genes analysed, split into only the *Wnt8a.1* genes in *E. antarctica*, the *Wnt8a.1* genes in all other species and then all other *Wnt* genes in all other species. *p-value:* *, < 0.05.

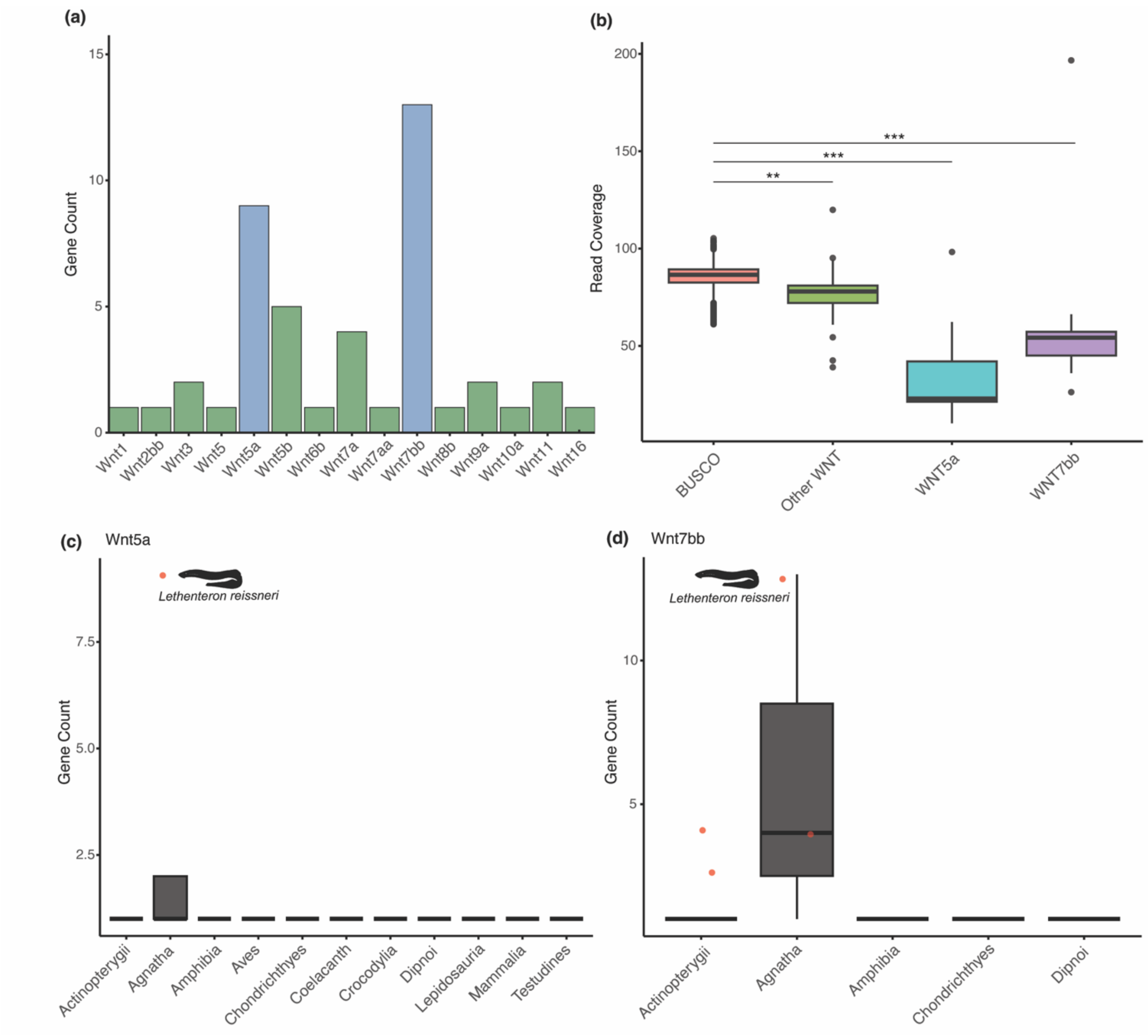


**Fig. S3. *Lethenteron reissneri Wnt5a* and *Wnt7bb* expansions. (a)** Gene copy number per *Wnt* subfamily in *L. reissneri* showing that the expansion of *Wnt* genes in this species is mostly due to *Wnt5a* and *Wnt7bb*. **(b)** A comparison of raw genomic read coverage between BUSCO genes, other *Wnt* genes (excluding *Wnt5a* and *Wnt7bb* copies), and only the *Wnt5a* or *Wnt7bb* genes in *L. reissneri*. *p-value:* **, < 0.01; ***, < 0.001. **(c)** The distribution of *Wnt5a* genes across all vertebrate species analysed, split by class. The species with the highest copy number of this gene (*L. reissneri*) is labelled. **(d)** The distribution of *Wnt7bb* genes across all vertebrate species analysed, split by class. The species with the highest copy number of this gene (*L. reissneri*) is labelled.

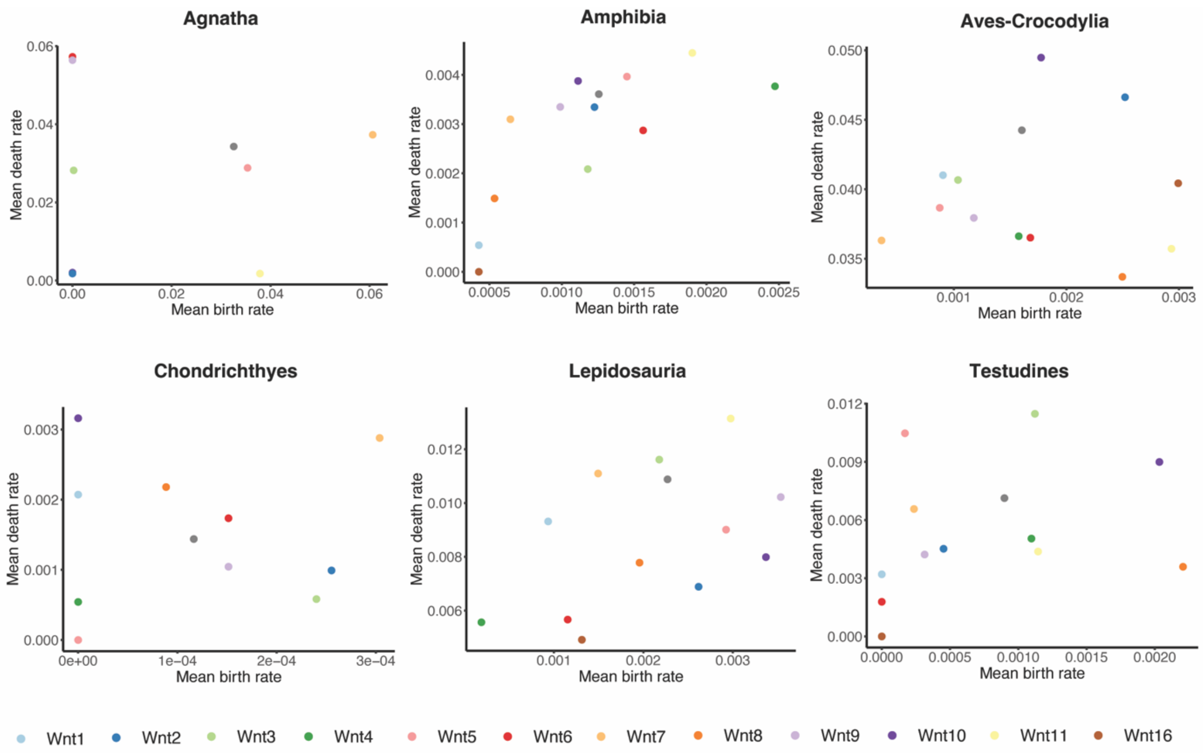


**Fig. S4. *Wnt* gene birth and death rates across vertebrate classes.** Mean per-gene subfamily death rate against mean birth rate plotted for the vertebrate classes, Agnatha, Amphibia, Aves-Crocodylia, Chondrichthyes, Lepidosauria and Testudines. Dots are coloured by *Wnt* gene subfamily. Note that the plots for Actinopterygii and Mammalia can be found in the main text.

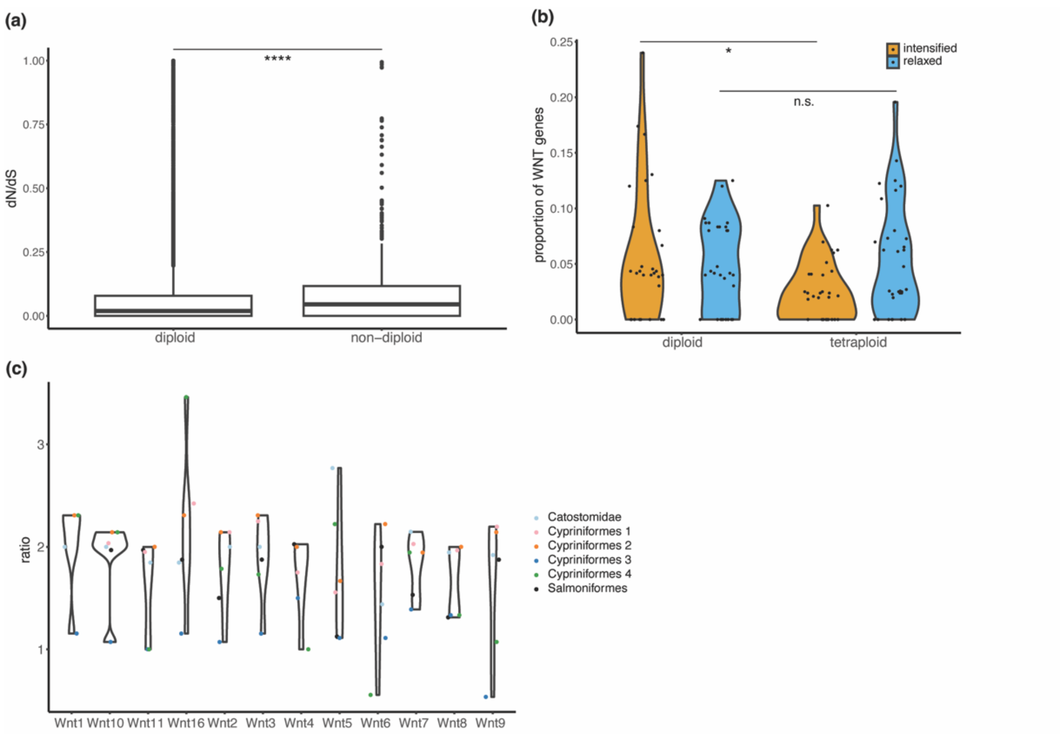


**Fig. S5. The fate of *Wnt* genes after whole-genome duplication (WGD) in ray-finned fishes. (a)** Boxplots showing the dN/dS values of *Wnt* genes in diploid and non-diploid species in Actinopterygii. **(b)** Violin plots of the proportion of *Wnt* genes under either relaxed or intensified selection in diploid and non-diploid species in Actinopterygii. **(c)** Violin plots of the mean ratio between the copy numbers of tetraploid species and closely related diploid species, per *Wnt* gene clade. Significance in (a) and (b) detected using a Wilcoxon signed-rank test. *p-value*: *, < 0.05; ****, < 0.0001.

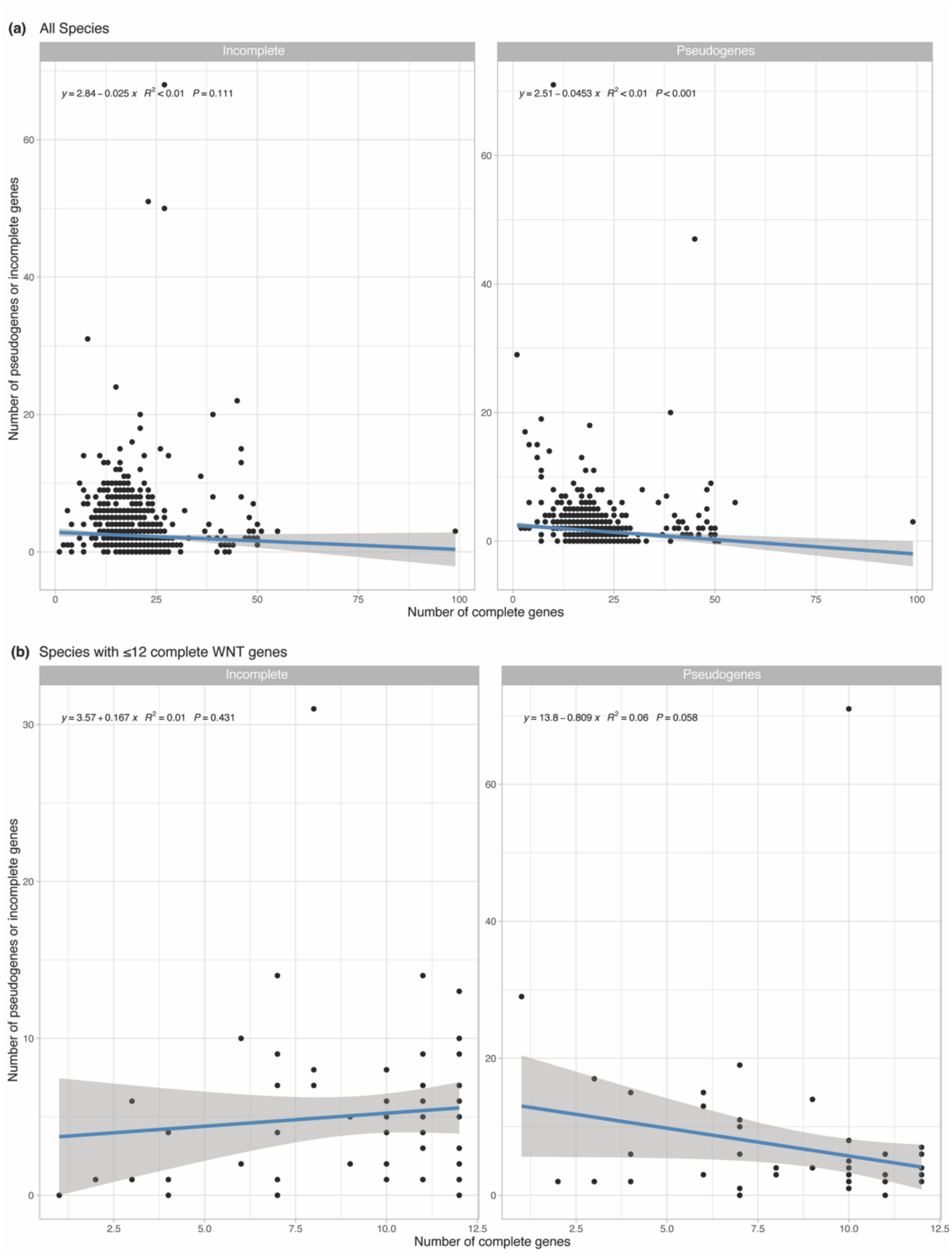


**Fig. S6. Relationship between number of complete *Wnt* genes and incomplete genes or pseudogenes. (a)** Correlation between number of complete *Wnt* genes and number of incomplete genes or pseudogenes for all species in this study. Equation, $R^2$ value and p-value were calculated for the linear regression (blue line). Note that there is a very weak negative correlation between the number of complete genes and pseudogenes but not incomplete genes. **(b)** Correlation between number of complete *Wnt* genes and number of incomplete genes or pseudogenes for species in this study which had fewer than 12 complete *Wnt* genes. Note that there is no correlation between the number of complete genes and pseudogenes but not incomplete genes. Equation, $R^2$ value and p-value were calculated for the linear regression (blue line).

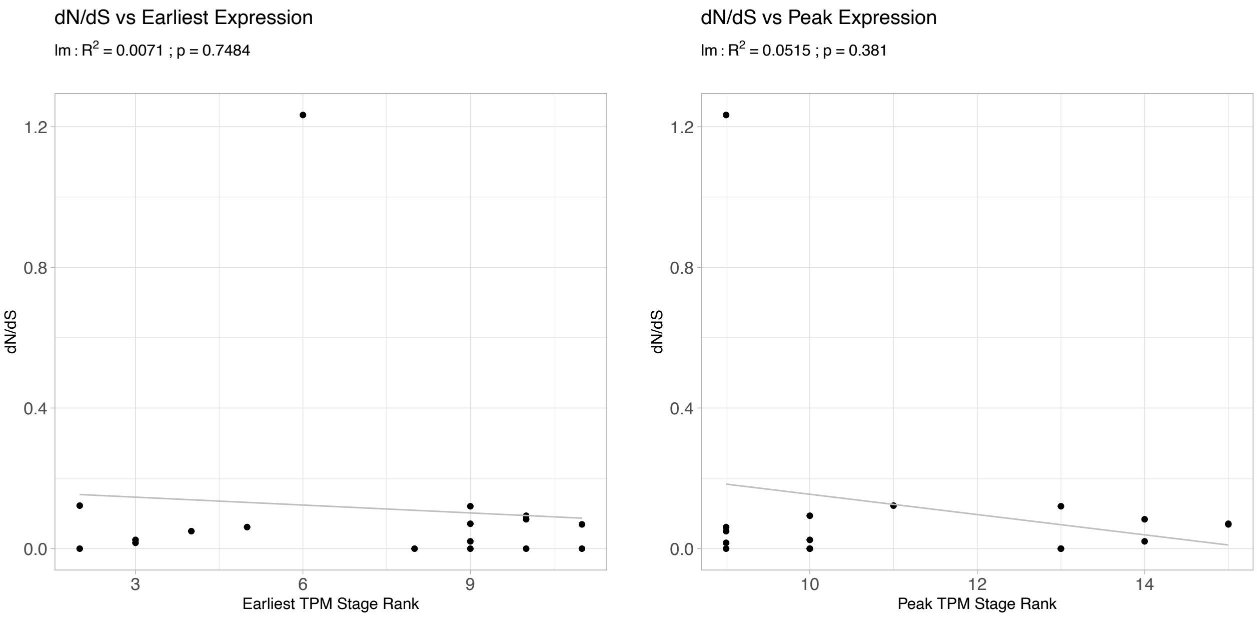


**Fig. S7. Relationship between gene expression and selective pressures acting on *Wnt* genes in *Danio rerio.*** Plots of dN/dS values of *Wnt* genes against the life stage (as a numerical rank) at which this gene showed its earliest (left) and peak (right) expression [as transcripts per million (TPM)]. Black line represents a linear regression, with the $R^2$ and p-values given above each plot.

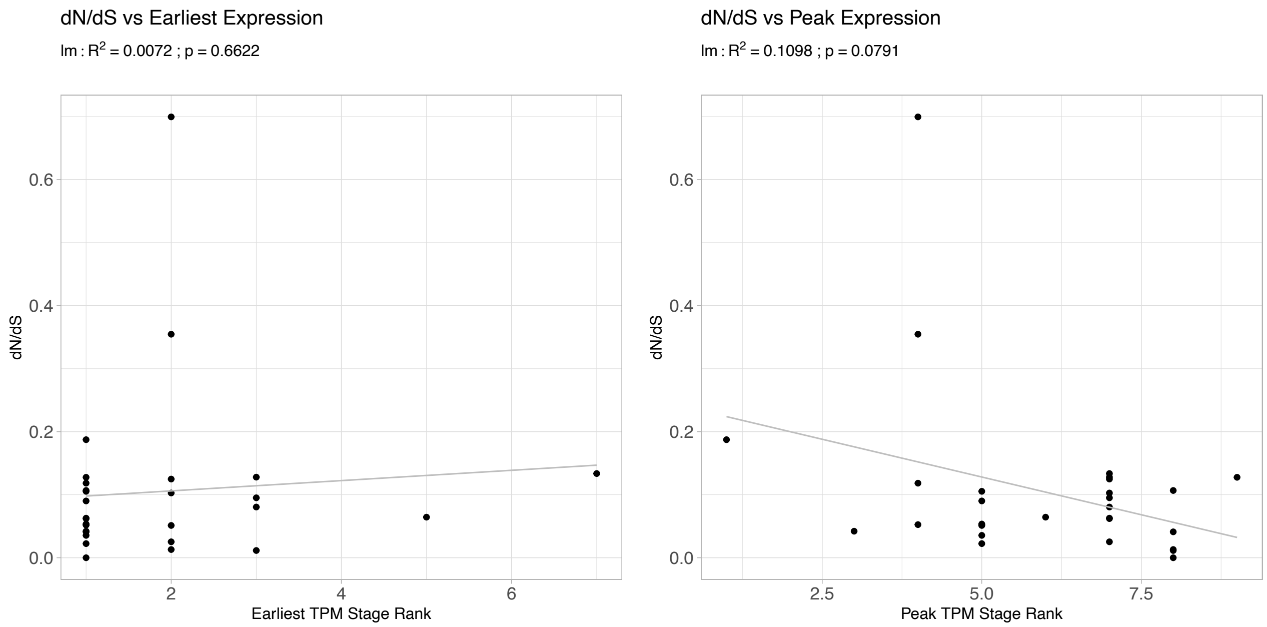


**Fig. S8. Relationship between gene expression and selective pressures acting on *Wnt* genes in *Xenopus laevis.*** Plots of dN/dS values of *Wnt* genes against the life stage (as a numerical rank) at which this gene showed its earliest (left) and peak (right) expression [as transcripts per million (TPM)]. Black line represents a linear regression, with the $R^2$ and p-values given above each plot.

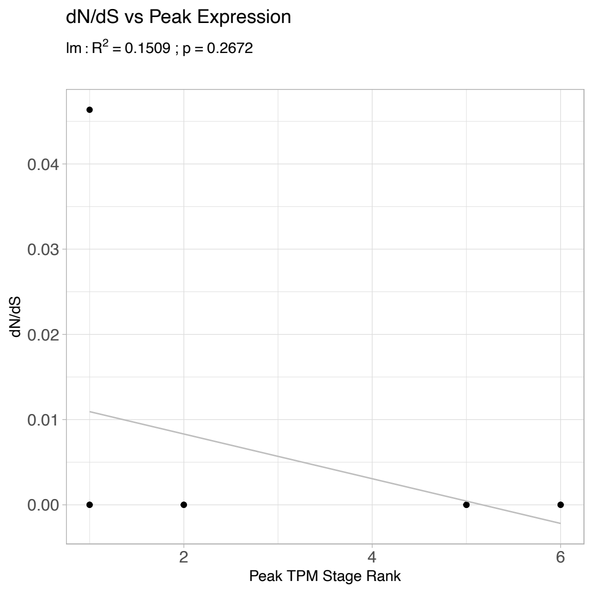


**Fig. S9. Relationship between gene expression and selective pressures acting on *Wnt* genes in *Mus musculus.*** Plots of dN/dS values of *Wnt* genes against the life stage (as a numerical rank) at which this gene showed its peak expression [as transcripts per million (TPM)]. Black line represents a linear regression, with the $R^2$ and p-values given above each plot.
